# Variation in phenotypic plasticity of metabolic and performance traits along a latitudinal gradient in Woodland Strawberry

**DOI:** 10.64898/2025.12.18.695113

**Authors:** José E. Pérez-Martín, Femke Batsleer, Martijn L. Vandegehuchte, Ivan M. De-La Cruz, Carolina Diller, José F. Sánchez-Sevilla, Anne Muola, Johan A. Stenberg, Timo Hytönen, Dries Bonte, David Posé, Sonia Osorio

## Abstract

Phenotypic plasticity and local adaptation jointly determine plant responses to environmental variation, yet their relative contributions and evolutionary interactions remain poorly understood. We grew fifteen woodland strawberry (*Fragaria vesca*) genotypes from a European latitudinal gradient in four common gardens for two years, quantifying primary metabolites and biomass, reproduction and herbivore damage-related traits. Considering all measured traits combined, environment explained the largest variance (30%), followed by genotype-by-environment interactions (18%) and genotype effects (9%), indicating substantial heritable plasticity. However, plasticity was not uniform: northern genotypes showed greater plasticity in stress metabolites like succinic acid but more canalized growth traits, whereas southern genotypes maintained constitutively high protective metabolite levels. Latitude explained 4.4% of plasticity profile variation. Phenotypic variation was consistently better explained by long-term temperature than precipitation at the site of origin across all trait groups under study. Our results indicate that plasticity itself is an evolved, trait-specific characteristic shaped by climatic gradients, challenging simple hierarchies of trait lability.

**Significance Statement:** We studied wild strawberries from across Europe and discovered that populations from different regions have evolved distinct adaptive strategies to cope with their local cli-mates. These differences highlight the remarkable diversity of plant solutions to environmental challenges. Understanding these natural adaptations could help us breed strawberries or other plant organisms better suited to future climate conditions.

## Introduction

Climate change represents an unprecedented global challenge, disrupting temperature and precipitation patterns, altering seasonal cues such as photoperiod-temperature mismatches, and increasing the frequency and intensity of extreme events such as heatwaves, droughts, hurricanes, and floods. These perturbations cascade through plant function: elevated temperatures and COLJ levels affect photosynthesis, respiration and growth; irregular or unpredictable rainfall compromises water-use efficiency; and changing winter conditions due to altered temperature and precipitation regimes threaten overwintering survival, particularly at higher latitudes and elevations, and storms or flooding cause mechanical damage that can delay flowering and fruiting (Jentsch et al., 2009; Wenzel et al., 2016; Weiskopf et al., 2020; Flores et al., 2023). Because theses organism-level effects scale up to influence community composition, ecosystem services, and even macro-evolutionary trajectories (Traill et al., 2010; Gornish and Tylianakis, 2013), understanding how plants respond to environmental change remains a central challenge in ecology and evolutionary biology.

Two non-exclusive mechanisms underpin a plant’s capacity to cope with environmental change: local genetic adaptation and phenotypic plasticity, defined as the ability of a single genotype to express different phenotypes across environments. Phenotypes are shaped by genetic factors (G), the local environment (E), and genotype-by-environment interactions (G×E), which reflect differential responses of genotypes to environmental variation. Plasticity can provide rapid, reversible buffering against stress while directional selection reshapes mean trait values. Plasticity is itself heritable and subject to selection, especially in environments that are heterogeneous yet predictable enough for reliable cues (Bradshaw, 1965; Via and Lande, 1985; Schlichting, 1986). In fact, plasticity can evolve into a specialized adaptation to frequent, spatial variable, or temporally unpredictable conditions (Hollander et al., 2015; Snell-Rood and Ehlman, 2021).

Comparing distinct sites across broad spatial scales provides natural laboratories for exploring these concepts (Joschinski and Bonte, 2021). Moving poleward, mean temperature declines and annual thermal amplitude generally increases, while day-length becomes a highly predictable seasonal cue (Gaskell et al., 2022). European precipitation patterns broadly follow an east-west gradient, with coastal western regions receiving abundant rainfall and inland eastern areas remaining much drier (Fick and Hijmans, 2017).

Plants track these broad environmental clines through coordinated shifts in metabolism, morphology, physiology and life-history (Ackerly, 2003; Valladares et al., 2007; Gratani, 2014). Metabolic osmolytes such as proline and organic acids accumulate in cooler, wetter sites prone to frost or waterlogging (Bartels and Sunkar, 2005; Planchet and Limami, 2015), while traits like leaf area and root-to-shoot ratio often canalize around local optima once construction costs outweigh further plasticity benefits (Bongers et al., 2017). Empirical syntheses associate high-latitude populations with relatively fixed phenological schedules driven by reliable photoperiods, but with broad biochemical capacity for cold acclimation. Low-latitude populations in thermally stable but rainfall-variable climates invest in drought-responsive metabolism while maintaining flexible water-use physiology (Reger et al., 2017; de Villemereuil et al., 2018; Schneider, 2022).

Trait classes are predicted to differ in plasticity: metabolic traits (e.g., sugar and amino acid concentrations) can shift within hours and are expected to be most labile (Murren et al., 2015; Butnariu and Bocso, 2022). However, recent meta-analyses reveal more complex patterns: morphological traits can sometimes exceed physiological traits such as photosynthetic capacity, stomatal conductance, and water-use efficiency in plasticity, challenging classic hierarchies (Acasuso-Rivero et al., 2019). Temporal dynamics are also critical: fast-responding traits track short-term weather fluctuations, while integrative demographic traits respond to long-term climate trends (De Lisle and Rowe, 2023).

Plant defense against herbivory represents another critical dimension of environmental response, where both constitutive and induced defenses can vary substantially among environments. Metabolites, both primary (proline, glucose) and secondary (phenolic compounds, alkaloids) like phenolic compounds and alkaloids provide chemical protection, while physical traits such as leaf toughness and trichome density offer mechanical barriers. The balance between constitutive and plastic defense strategies often reflects the predictability of herbivore pressure across environments, with populations from high-herbivory sites typically maintaining higher baseline defenses while retaining capacity for further induction when attacked (Agrawal, 2001; Karban and Baldwin, 2007).

*Fragaria vesca* (woodland strawberry) is an ideal model for trait-specific plasticity studies due to its broad ecological range (from temperate forests to alpine meadows across Europe, Asia, and North America), combined with high intraspecific genetic diversity and a fully annotated genome (Shulaev et al., 2010; Edger et al., 2018; Li et al., 2019; Zhou et al., 2023). Its ease of cultivation and clonal propagation have facilitated eco-evolutionary experiments documenting pronounced plastic shifts in leaf morphology and flowering time (Heide and Sønsteby, 2007; De Kort et al., 2020; De-la-Cruz et al., 2025b). Importantly, *F. vesca* constitutes the dominant ancestral sub-genome of cultivated octoploid strawberry *Fragaria* × *ananassa*, making mechanistic insights directly relevant to crop improvement (Edger et al., 2019).

Using a European germplasm collection of approximately 200 accessions that captures genome-wide and volatile diversity (Urrutia et al., 2023; Toivainen et al., 2024), we selected fifteen genotypes spanning a latitudinal gradient and cultivated them for two years in common garden trials at four contrasting sites (Gontrode, Belgium; Alnarp, southern Sweden; Ruissalo, southern Finland; and Kevo, northern Finland). With this design we tested the prediction that plasticity is highest in primary metabolic traits (sugars, amino and organic acids), moderate in physiological and morphological traits, and lowest in integrative demographic metrics such as biomass or fruit yield.

By quantifying metabolic, biomass, demographic, and herbivore damage-related traits across environments, we assess their variation in plasticity, identify trade-offs among trait classes, and elucidate how *F. vesca* reallocates resources via metabolic reprogramming versus structural change to cope with climate-relevant stresses (Comas et al., 2013; Joshi, 2023). We further explore how plasticity relates to genotypes’ evolutionary history and temporal scales of environmental variation they have experienced.

## Results

### Characterization of trait variation and covariation

Fifteen *Fragaria vesca* genotypes collected across a south-north latitudinal gradient in Europe (Figure 1) were grown in five common garden environments and analyzed for metabolic and performance traits. Metabolite profiling led to the identification of 36 primary metabolites (Table S2; raw data deposited in Zenodo: *it will be deposited upon acceptance*), which in general showed strong grouping according to the GARDEN_ID (Figure 2). Metabolites known to be involved in closely related biochemical pathways often exhibited similar abundance patterns across environments, resulting in their clustering. Six metabolite clusters emerged, primarily separating organic acid and amino acid pools. Plants grown at the Ruissalo location showed the lowest concentrations across most clusters suggesting a generally slower metabolism at that location. Of the 36 metabolites measured, only urea, fructose and glucose showed no significant association with any of the climate variables considered. The strongest climate signals were observed in a cluster mainly including organic acids (along with several amino acids). In addition, aspartic acid, glutamic acid, and pyroglutamic acid increased with precipitation (r = 0.56 to 0.77) but declined with mean, maximum, and minimum temperatures (r = -0.47 to -0.79). A second cluster, largely composed of nitrogen-related compounds, showed a similar pattern, proline and threonine increased with precipitation (r = 0.61 to 0.64) and decreased with maximum temperature (r = -0.63 to -0.65). In contrast, tryptophan, glycine, and pyruvic acid showed the opposite pattern, decreasing with rainfall (r = -0.49 to -0.82) and increasing with warmer temperatures (r = 0.32 to 0.80).

**Figure 1:**
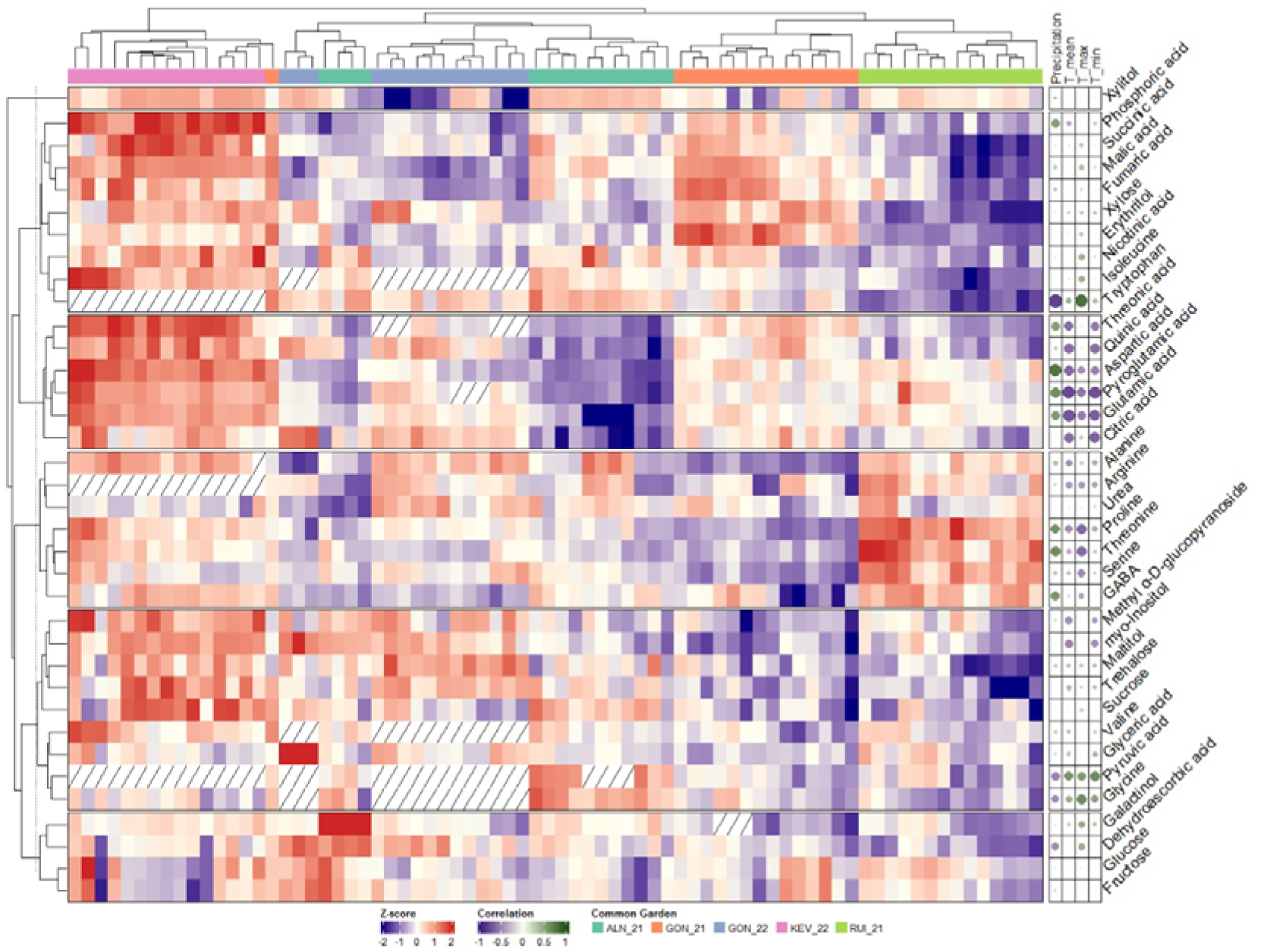
Clustered heatmap of the relative abundance of 36 primary metabolites (sugars, amino acids, organic acids, and others) in young leaves. Row clustering (metabolites) was performed using hierarchical clustering (Euclidean distance, complete linkage), divided into 6 clusters. The Pearson correlations with climatic variables are represented by dotted heatmap on the right side of the plot (metabolites vs. precipitation, mean, maximum, and minimum temperature).

**Figure 2:**
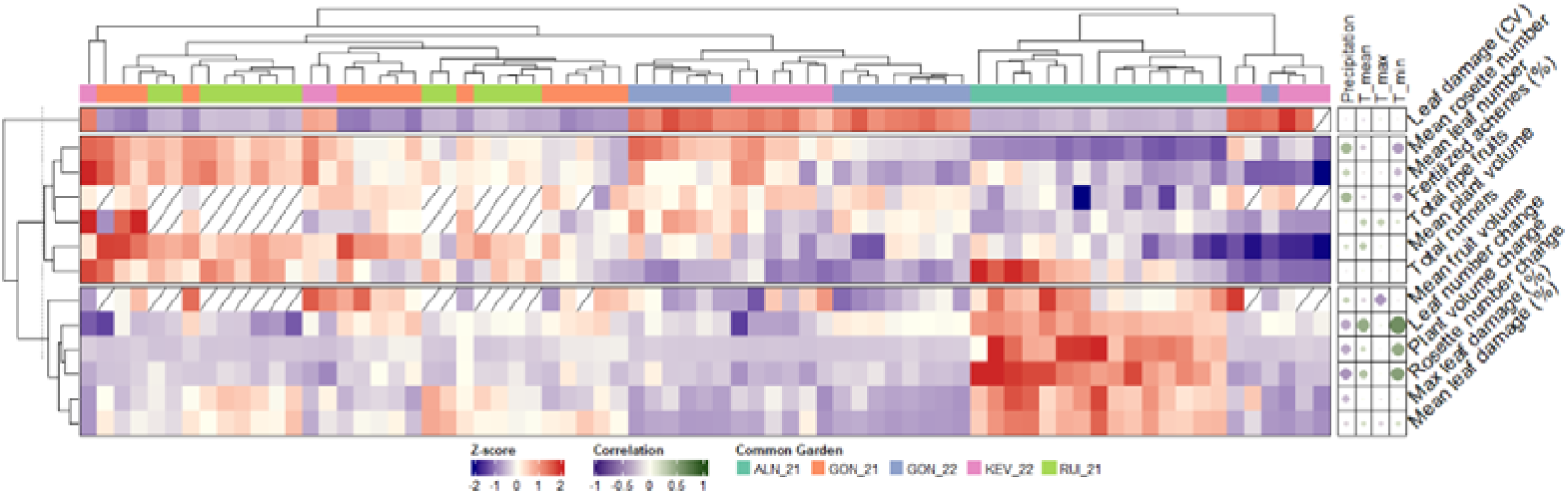
Clustered heatmap of the 13 measured performance-related traits (biomass, reproduction, and herbivore damage-related). Row clustering of traits was performed using hierarchical clustering (Euclidean distance, complete linkage), divided into 3 clusters. The Pearson correlations with climatic variables are represented by dotted heatmap on the right side of the plot (traits vs. precipitation, mean, maximum, and minimum temperature).

Plant performance-related traits (Table S3, raw data deposited in Zenodo: *it will be deposited upon acceptance*) also grouped by GARDEN_ID, although their associations with climate variables were generally weaker than those observed for metabolites (Figure 3). Several key traits, including mean plant volume, total number of runners, and mean leaf damage (%), showed no direct correlation with the measured annual climate variables. However, a clear geographic trend emerged, plants grown in the northern common gardens displayed reduced overall growth (e.g., plant volume change) and reproductive output (e.g., total ripe fruits). Year-to-year variation at the Gontrode location was particularly pronounced for herbivore damage-related traits. The strongest climatic signal was observed for biomass traits, where leaf and rosette change declined with increasing precipitation (r = -0.36 to -0.45) but increased with higher mean and minimum temperatures (r = 0.33 to 0.62).

**Figure 3:**
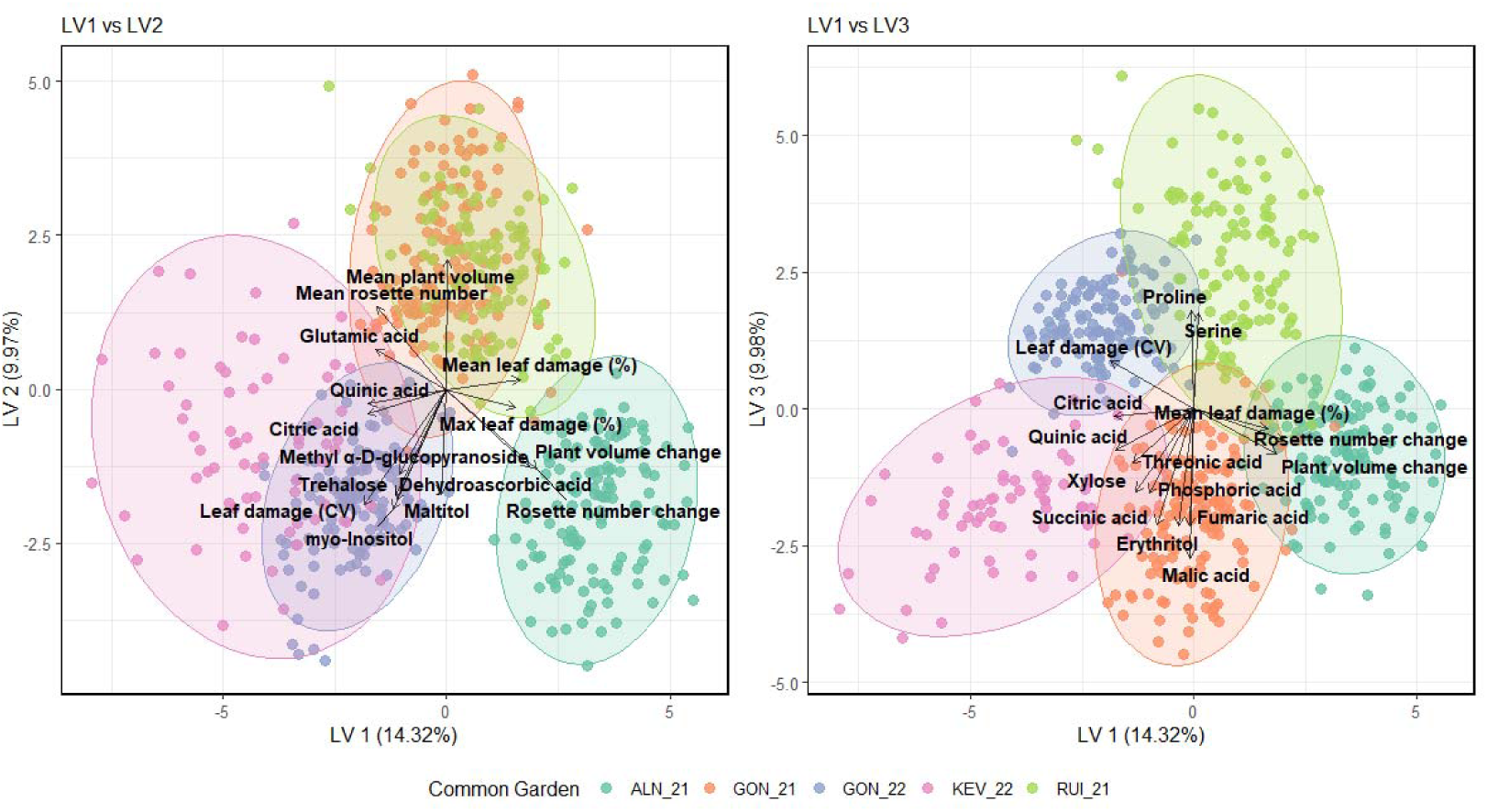
Partial Least Squares Discriminant Analysis (PLS-DA) biplots of combined performance-related and metabolic data. The analysis shows the discrimination of plants grown in the five different common garden environments (KEV_22, ALN_21, RUI_21, GON_21, GON_22). (Left) Biplot of the first two latent variables (LV1 vs. LV2). (Right) Biplot of the first and third latent variables (LV1 vs. LV3). In both plots, points represent individual samples, colored by their garden of origin, with ellipses indicating the 95% confidence interval for each group. Vectors represent the loadings of the 15 most influential variables, indicating their contribution to the separation along the axes.

Following this preliminary analysis, we integrated all datasets using Partial Least Squares Discriminant Analysis (PLS-DA) to explore phenotypic clustering among the common gardens (Figure 4). The three main latent variables (LV1: 14.37%, LV2: 9.97%, LV3: 9.98%) contributed relatively equal to the model, highlighting their importance in explaining variability within the dataset. The PLS-DA plot clearly separated samples from KEV_22 from those of ALN_21, RUI_21, and GON_21 along LV1. KEV_22 experienced the most extreme climatic conditions, and even though RUI_21 had environmental parameters most similar to KEV_22 of the other southern gardens, their trait profiles diverged strongly along LV1. Further separation between GON_21 and GON_22 were driven by LV2, while LV3 distinguished ALN_21 from RUI_21. Notably, the separation between GON_21 and GON_22, despite their shared location, highlights the strong influence of short-term environmental variation on plant performance, with 2021 being an exceptionally wet year at that site (Journée et al., 2023).

**Figure 4:**
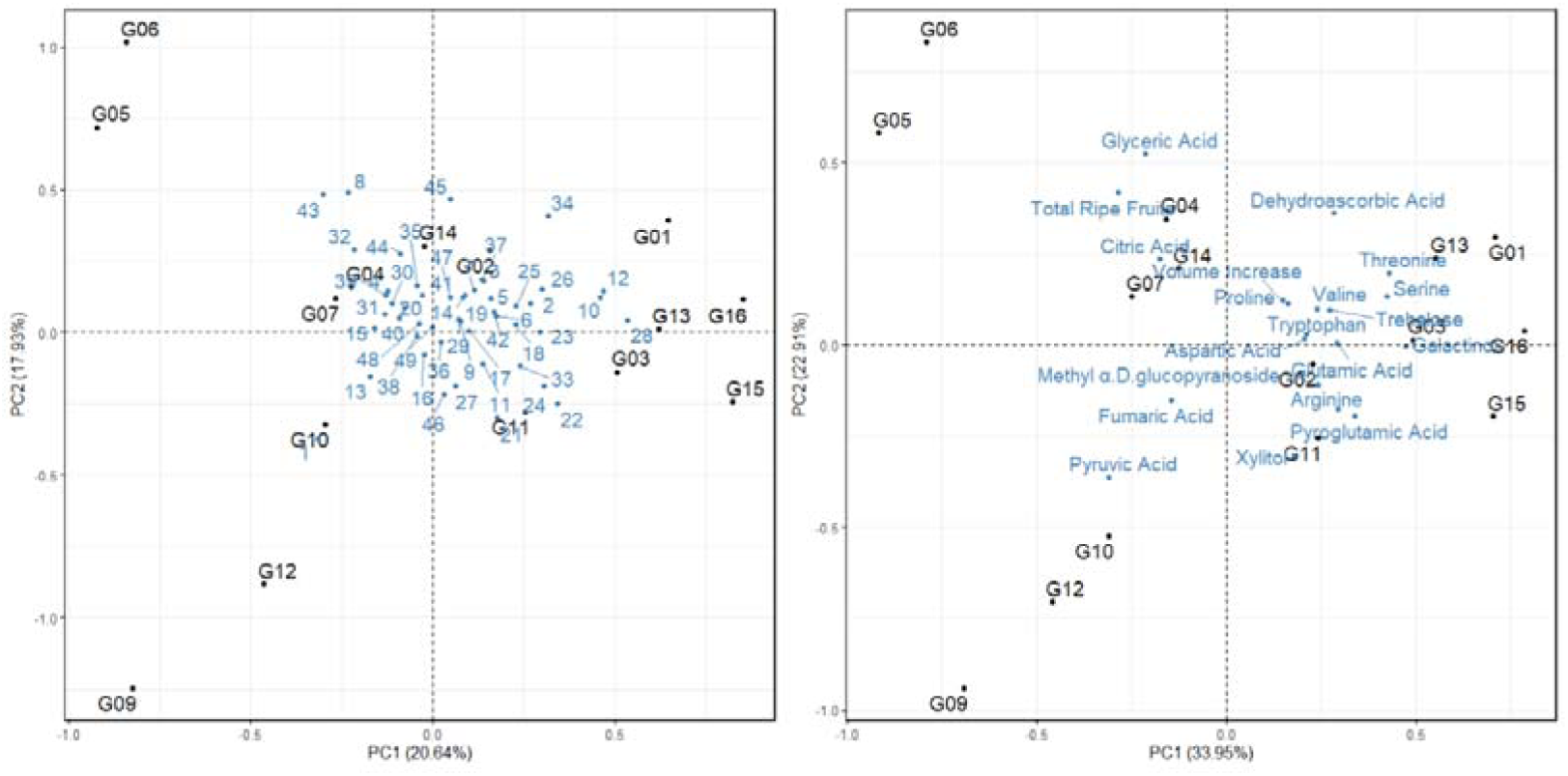
PCA of ICCs quantifying genotype-level phenotypic plasticity. (A) Biplot using all measured traits, each labelled by a trait number. (B) Biplot using the 20 traits with the highest contributions, labelled by the same trait numbers. See Table S4 for the full mapping of trait numbers to trait names.

The variable loadings, visualized as vectors in the biplots, reveal the key drivers of this separation (Figure 4). Consistent with our previous observations, the clear clustering of KEV_22 was strongly influenced by higher values of several important organic acids, such as citric acid, and by lower leaf damage percentage, suggesting these metabolites may contribute directly to herbivore resistance rather than indicate a trade-off with it. The separation between GON_21 and GON_22 appeared to be driven by specific organic acids and sugars, while performance-related traits contributed less to this separation. Finally, the separation of RUI_21 was principally associated with amino acids, including proline, which plays a key role in stress response.

PERMANOVA analysis indicated that the environment (GARDEN_ID) was the principal factor explaining phenotypic variation, accounting for 30.38% of the total variability when entered first in the model, while genotype explained 9.40% and the G×E interaction for 17.89% (Table 1). To address the sequential sum-of-squares nature of PERMANOVA, we ran an alternative model in which genotype was entered first and environment second: environment then explained 29.41% and genotype 10.37%. In both cases, environment remained the strongest predictor, and the minimal difference in explained variance (< 1% for both terms) confirms the robustness of these results regardless of model order.

**Table 1:** Results of Permutational Multivariate Analysis of Variance (PERMANOVA) testing the effects of environment (GARDEN_ID), genotype, and their interaction on multivariate phenotypic traits. Terms were added sequentially (Type I sum of squares) in the order GARDEN_ID, Genotype, GARDEN_ID × Genotype. Df = numerator degrees of freedom; ddf = denominator degrees of freedom; SumOfSqs = sum of squares; R² = proportion of total variance explained (SumOfSqs/Total SumOfSqs); F (ndf, ddf) = FLJstatistic with degrees of freedom; Pr(>F) = PLJvalue based on 9999 free permutations. Asterisks (***) denote p < 10LJLJ. An alternative model with Genotype entered before GARDEN_ID yielded comparable R² values (Genotype = 10.4%, GARDEN_ID = 29.4%), confirming the robustness of the environment’s predominant effect despite model order.

|  | **Df** | **SumOfSqs** | **R<sup>2</sup>** | **F (ndf, ddf)** | **Pr(>F)** |
| --- | --- | --- | --- | --- | --- |
| **GARDEN_ID** | 4 | 7411.7 | 30.38 | 86.35 (4, 509) | $< 10^{-4}$ *** |
| **Genotype** | 14 | 2406.2 | 9.40 | 8.07 (14, 509) | $< 10^{-4}$ *** |
| **GARDEN_ID:Genotype** | 53 | 4587.9 | 17.89 | 4.05 (53, 509) | $< 10^{-4}$ *** |
| **Residual** | 509 | 10922.2 | 42.33 |  |  |
| **Total** | 580 | 25328.0 | 100.00 |  |  |

Phenotypic variation thus primarily reflects environmental effects, with genotype and genotype-by-environment interaction contributing substantially, revealing both strong plasticity and a significant genetic basis that could be subject to evolutionary selection. These results establish that immediate growing conditions (GARDEN_ID) dominate the phenotypic landscape at the among-garden scale, independent of genotype identity.

To further partition variance between immediate growing conditions (GARDEN_ID) and phenotypic signatures associated with each genotype’s climatic origin (extracted from its collection coordinates), we performed sequential db-RDA analysis with conditional modeling. This approach tests whether genotypes retain adaptive trait signatures from their native climate independent of current growing environment. This analysis revealed that GARDEN_ID dominated all trait categories, explaining 31.53% of variance in metabolic traits, 35.30% in biomass, 23.06% in reproduction, and 47.81% in herbivore damage (Table 2). After conditioning on GARDEN_ID effects, Annual Mean Temperature (BIO_1) at genotype origin consistently outperformed precipitation (BIO_12) as a predictor of additional trait variance.

**Table 2:** Results from distance-based Redundancy Analysis (db-RDA) with sequential conditioning, partitioning phenotypic variance according to growing ment (GARDEN_ID) and bioclimatic origin covariates. For each trait group, two separate conditional models were constructed where the bioclimatic (Annual Mean Temperature, BIO_1; or Annual Precipitation, BIO_12) was tested with GARDEN_ID effects removed [Condition(GARDEN_ID)], g the independent contribution of native climate to phenotypic variation. R² (%) Garden Environment represents adjusted variance explained by EN_ID alone; R² (%) Climatic Covariate represents adjusted variance explained by the bioclimatic variable after conditioning on GARDEN_ID effects. tics and p-values are from permutation tests with 9999 free permutations. Asterisks (***) denote p < 10ll. Non-significant p-values for herbivore traits indicate no detectable signal of local adaptation to native climate for these traits despite strong GARDEN_ID effects.

| Trait Group | Climatic Driver | R<sup>2</sup> (%) (Climatic Covariate) | R<sup>2</sup> (%) (Garden Environment) | F-statistic | P-value |
| --- | --- | --- | --- | --- | --- |
| Metabolic | Temperature (BIO1) | 0.81 | 31.53 | 7.91 | $< 10^{-4}$ *** |
| | Precipitation (BIO12) | 0.58 | 31.53 | 5.89 | $< 10^{-4}$ *** |
| Biomass | Temperature (BIO1) | 2.52 | 35.30 | 24.33 | $< 10^{-4}$ *** |
| | Precipitation (BIO12) | 1.67 | 35.30 | 16.29 | $< 10^{-4}$ *** |
| Reproduction | Temperature (BIO1) | 1.78 | 23.06 | 14.66 | $< 10^{-4}$ *** |
| | Precipitation (BIO12) | 0.97 | 23.06 | 8.34 | $< 10^{-4}$ *** |
| Herbivore damage | Temperature (BIO1) | -0.05 | 47.81 | 0.49 | 0.578 |
|  | Precipitation (BIO12) | 0.00 | 47.81 | 0.99 | 0.337 |

For metabolic traits, temperature contributed an additional 0.81% of variance (p < 10^-4^), while biomass traits showed the strongest temperature signal at 2.52% (p < 10^-4^). Reproduction traits exhibited an intermediate response with 1.78% additional variance explained by temperature (p < 10^-4^). Precipitation contributed substantially less across trait groups, never exceeding 1.67%. In contrast, herbivore damage traits showed a fundamentally different pattern. Although GARDEN_ID dominated (47.81%, p < 10^-4^), neither temperature (R² = -0.05%, p = 0.578) nor precipitation (R² = 0.00%, p = 0.337) contributed significant additional variance, indicating that herbivore damage is governed almost exclusively by plastic responses with no detectable signal of local adaptation to native climate. These results suggest a hierarchical structure: immediate environment effects dominate all traits; temperature-mediated adaptation is strongest for biomass, moderate for reproduction, and weak for metabolism but absent for herbivore damage; and precipitation plays only a subordinate role throughout.

### Patterns in phenotypic plasticity

Intraclass Correlation Coefficients (ICCs) were calculated for each trait to quantify the consistency of trait expression across GARDEN_IDs for each genotype. PCA of the full ICC matrix (Figure 5A) revealed that genotypes from mid-latitudinal origins (e.g., G05, G06, G09) exhibited the most divergent plasticity profiles among the cohort, characterized by unique combinations of highly plastic and highly canalized traits. In contrast, both southernmost and northernmost genotypes clustered more tightly near the origin, indicating more similar and intermediate plasticity strategies across traits. A targeted PCA on the 20 traits contributing most strongly to PC1 (Figure 5B) produced an equivalent separation, confirming that variation in these key traits drives the observed genotype divergence.

**Figure 5:**
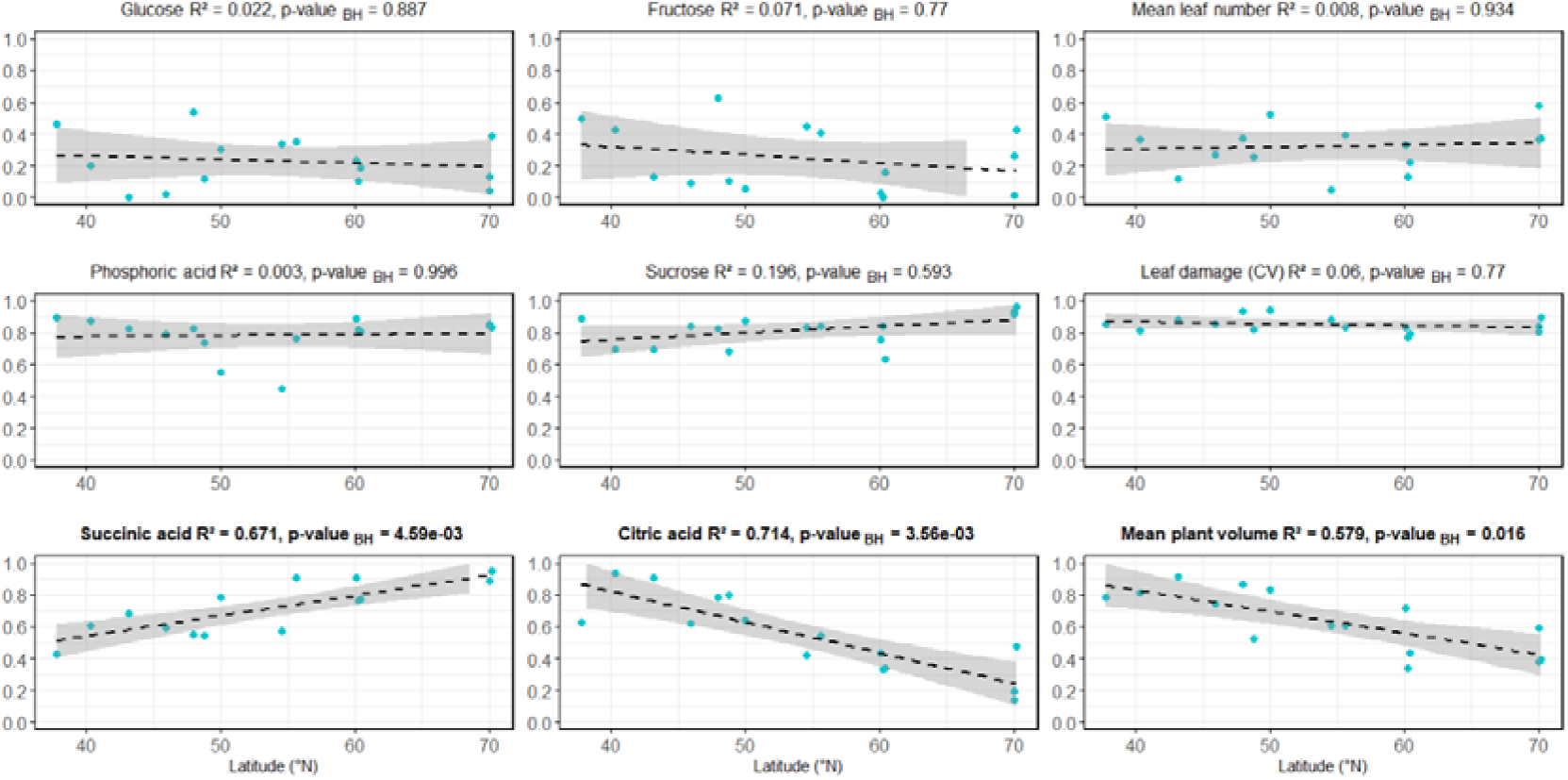
Scatter plots showing the relationship between latitude and ICCs for nine representative traits, with BH-adjusted p-values. Traits were selected to illustrate three distinct plasticity patterns: consistently low plasticity across genotypes (first row: glucose, fructose, mean leaf number), consistently high plasticity across genotypes (second row: phosphoric acid, sucrose, leaf damage (CV)), and latitude-dependent plasticity with significant latitudinal trends (last row: succinic acid, citric acid, mean plant volume). Relationships between latitude and ICCs for all 49 individual traits are provided in Figure S1.

To formally assess whether overall plasticity strategies were structured by geography, we db-RDA on the full ICC matrix. The analysis revealed a significant relationship between genotype plasticity profile and latitude of origin (F = 1.64; R²Adj = 0.044; p = 0.044), indicating that latitude explained 4.4% of the total variation. This suggests that selection along the latitudinal gradient has played a modest but significant role in shaping the genotypes’ strategies for responding to environmental change.

Correlations between latitudinal origin of each genotype and ICC values for individual traits highlighted notable variation in plasticity response. After adjusting p-values for multiple comparisons using the Benjamini-Hochberg (BH) procedure, several contrasting trends emerged (Figure 6, Figure S1). For instance, plasticity in succinic acid increased significantly with latitude (r = 0.82; p-valueBH = 2.99×10DD), indicating greater environmental sensitivity in northern genotypes. In contrast, citric acid (r = -0.84; p-valueBH = 1.86×10D³) and mean plant volume (r = -0.87; p-valueBH = 1.51×10D³) showed significantly reduced plasticity at higher latitudes, suggesting more stable trait expression.

**Figure 6:**
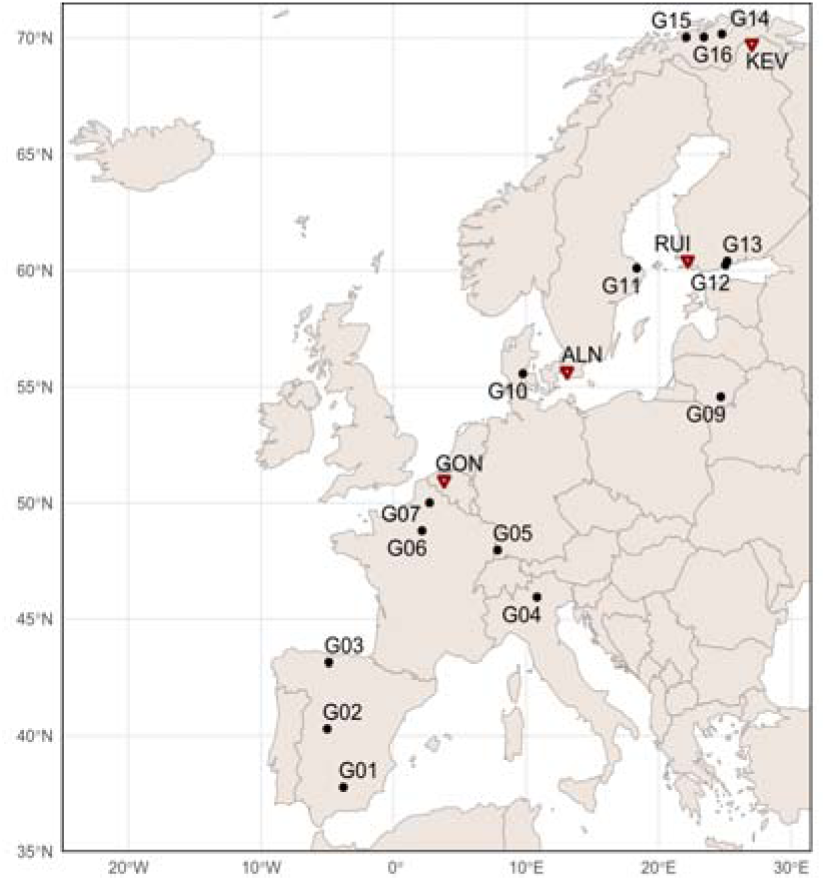
Map of Europe showing the origin locations of the fifteen selected *Fragaria vesca* genotypes (circles) and the common garden sites (red triangles). The genotypes were chosen to represent a south-north latitudinal gradient, and their numbering (G01 to G16, excluding G08) corresponds to the progressively increasing latitude of their origin.

These findings reinforce our earlier observations and support the hypothesis of complex trait-specific regulation, where responses are shaped by genotype-by-environment interactions. As a result, generalizing metabolic or performance-related responses across latitudinal clines is not feasible, underscoring the variability in environmental adaptation across both traits and genotypes.

## Discussion

In woodland strawberry, phenotypic variation is shaped by a complex interplay between phenotypic plasticity and local adaptation. Across our four European common gardens, short-term weather, climatological, and seasonal differences explained more trait variance than genotype, showing that where a plant grows often outweighs what genotype it is in determining the phenotype. However, our results challenge the notion of a simple plasticity hierarchy. Instead, we found trait-specific patterns of plasticity that varied independently across metabolic and performance-related traits. Some of them, particularly within primary metabolism, exhibited a pronounced dichotomy: several metabolites such as malic acid and phosphoric acid displayed high plasticity across all populations, whereas others, such as glucose and fructose, remained consistently canalized across genotypes. Moreover, for many other traits spanning from metabolic (e.g., pyruvic acid) to biomass (e.g., plant volume), the degree of plasticity itself was strongly influenced by the genotypes’ climate of origin. Most notably, demographic traits directly linked to fitness, such as fruit volume, defied expectations of canalization. Rather than being stable, they exhibited genotype-specific strategies, ranging from highly responsive in some genotypes to nearly fixed in others. These divergent responses highlight the prevalence of genotype-by-environment interactions, which accounted for approximately 18% of total variance. This interaction created contrasting plasticity profiles among genotypes rather than universally “plastic” or “rigid” genotypes. Together, these findings reveal that adaptation involves a complex interplay of flexible and constrained traits, often shaped by evolutionary history, offering valuable insights for both biological conservation and the breeding of climate-resilient crops.

### Common gardens demonstrate the importance of environmental drivers for phenotypic variation

The common garden design highlighted the dominant role of environmental conditions in shaping phenotypic variation. Most of the 36 quantified metabolites showed a strong association with local climatological conditions, including year-to-year variation, and were organized in different response clusters. Several organic acids and amino acids, such as aspartic acid, glutamic acid, and pyroglutamic acid, were positively correlated with precipitation and negatively with temperature, suggesting a role in adaptation to water availability and temperature stress (Bartels and Sunkar, 2005; Planchet and Limami, 2015). Other metabolites, including threonic acid, quinic acid, and citric acid followed similar but weaker correlations, suggesting a common regulatory network (Planchet and Limami, 2015; Tahjib-Ul-arif et al., 2021). The strong correlations between proline, threonine, and precipitation/temperature further support their role in stress responses (Kaur and Asthir, 2015; Planchet and Limami, 2015). On the other hand, tryptophan, pyruvic acid, and glycine showed the opposite behavior, negatively correlated with precipitation and positively with temperature, suggesting an alternative adaptive strategy focused on metabolic reconfiguration. This may involve increased production of signaling molecules derived from tryptophan (Corpas et al., 2021), enhanced osmoprotection via glycine-derived compounds like glycine betaine (Sakamoto and Murata, 2002), and adjustment to central energy pathways involving pyruvic acid as key node (Panchal et al., 2021). Only three metabolites showed weak correlations with environmental parameters, underscoring the pervasive influence of local conditions on plant metabolism.

Performance-related traits were also shaped by local environmental conditions, though their response patterns were more complex than those of metabolites. These traits did not follow a simple hierarchy of high/low plasticity; instead, their variation reflects highly genotype-specific strategies. For instance, the general decline in growth and reproductive output observed in northern common gardens (Figure 3) supports the idea that performance is reduced under less favorable conditions (Vergeer and Kunin, 2013). However, the degree of this reduction varied widely among genotypes, as shown by our ICC analysis. Similarly, large year-to-year variation in herbivore damage-related traits in specific populations highlights how short-term environmental fluctuations can trigger strong responses, but only in genotypes predisposed to such plasticity (Ibáñez et al., 2013).

### Trait-Specific Evolution of Phenotypic Plasticity

The heterogeneous plasticity patterns we observed across traits reflect complex regulatory mechanisms shaped by local adaptation (Schlichting, 1986; Schneider, 2022). Rather than supporting a universal hierarchy in which metabolic traits are more plastic than performance-related ones, our results reveal at least three distinct adaptive strategies (Figure 6).

First, some traits exhibited consistently low plasticity (low ICC values in Fig. 6) across all genotypes (e.g., glucose, fructose) suggesting strong stabilizing selection. Despite being part of primary metabolism, these metabolites are tightly regulated due to their central role in energy homeostasis, making their stability an adaptive feature rather than a constraint (Granot et al., 2013). The low ICC values for these traits indicate that each genotype maintains a characteristic metabolic profile across environments, pointing to fundamental genetic and developmental constraints that preserve trait stability (Takahashi, 2019; Snell-Rood and Ehlman, 2021).

Second, other traits showed consistently high plasticity (high ICC values in fig. 6), indicating that for certain functions, responsiveness is a universally advantageous strategy. This strategy is exemplified by metabolites like phosphoric acid, sucrose, and *myo*-inositol, that enable rapid adjustments to fluctuating nutrient availability and unpredictable stresses (Eckardt, 2010; Plaxton and Tran, 2011; Ruan et al., 2013; Weber et al., 2020). High plasticity in leaf damage metrics likely reflects both variable herbivore/pathogen pressure across gardens and genotypeDspecific tolerance mechanisms, rather than defense alone.

The third pattern is latitude-dependent plasticity. In northern genotypes, traits such as succinic acid showed significantly greater plasticity, whereas others, like citric acid and plant volume, were more canalized. For plant volume, this can be explained by the consistent compact growth form found in northern genotypes, even then transplanted southwards (De-la-Cruz et al., 2025a), and adaptive strategy in cold regions preserving heat (Neuner et al., 2000; Cranston et al., 2015). This demonstrates that plasticity itself is a locally adapted trait (Kawecki and Ebert, 2004), where selection has favored different degrees of environmental sensitivity in different native environments (Bradshaw, 1965; Reger et al., 2017). This trait-genotype-specific evolution of plasticity explains why plants stabilize some pathways while maintaining flexibility in others, preventing any single rule for predicting plastic responses.

An intriguing pattern emerged from our PCA of plasticity profiles: genotypes from mid-latitudinal origins exhibited the most divergent combinations of plastic and canalized traits, whereas both southern and northern genotypes converged on more similar strategies. This suggests that populations at intermediate latitudes may experience greater environmental heterogeneity or occupy transitional ecotones where multiple adaptive strategies remain viable. In contrast, extreme environments at both latitudinal ends may impose stronger directional selection, favoring convergence on optimal trait combinations tailored to predictable abiotic constraints. This pattern warrants further investigation into the role of climatic variability and habitat heterogeneity in shaping the evolution of plasticity architectures across geographic gradients.

### Latitude-Dependent Plasticity and its Adaptive Significance

The plasticity of certain traits varied predictably with the latitude of origin, suggesting that plasticity itself is a key adaptive trait subject to selection. Our results reveal opposing patterns across traits, which can be interpreted as distinct evolutionary strategies for coping with predictable versus unpredictable environmental stress.

A notable example is succinic acid, a key intermediate in the Krebs cycle and an important metabolite in stress tolerance and cellular energy metabolism (Khan et al., 2020), whose plasticity significantly increased in high-latitude genotypes. This might seem counterintuitive but aligns with adaptation to different stress patterns. In the low-latitude, high-elevation mountain climates of southern Europe, plants face a wide range of unpredictable daily and seasonal stresses, including intense heat, summer drought, and freezing winter temperatures. Under such variable conditions, maintaining constitutively high levels of protective metabolites like succinic acid (i.e., low plasticity) may represent a bet-hedging strategy, ensuring constant readiness without the need for energetically costly adjustments (Khan et al., 2020). High-latitude climates are characterized by greater environmental unpredictability, with frequent temperature fluctuations that easily cross the freezing point (0°C) and photosynthetic point (4°C), particularly during spring and autumn transitions (Nievola et al., 2017). Under such highly variable conditions, high plasticity in stress-response metabolites like succinic acid enables dynamic adjustment of cellular energy metabolism, with an increased production during cold acclimation and decreased levels during the growing season, offering a more energy-efficient strategy (Panchal et al., 2021; Kiliç, 2023). In contrast, southern European populations experience more seasonally predictable stress patterns. Under these stable conditions, maintaining constitutively high levels of protective metabolites (which is, exhibiting low plasticity), represents a more efficient strategy, ensuring constant readiness without requiring energetically costly metabolic adjustments (Khan et al., 2020).

In contrast, plasticity in citric acid and plant volume decreased in high-latitude genotypes. Like succinic acid, citric acid is a central metabolite in the Krebs cycle that plays a key role in energy regulation (Wang et al., 2017). The short, intense growing seasons in northern regions impose strong selective pressure on northern genotypes to maximize metabolic efficiency and survival. For citric acid, we hypothesize that this canalization reflects streamlining of metabolic energy production. For plant volume, the low plasticity is probably related to a compact growth form. This phenotype, which persists in northern genotypes even when transplanted to warmer climates (De-la-Cruz et al., 2025a), is a known adaptation to cold environments that minimizes heat exchange and maximizes solar warming (Körner et al., 2023).

For several drought-responsive amino acids and related metabolites (glutamic acid, arginine, methyl-α-D-glucopyranoside), we did not detect significant latitudinal trends in plasticity (pBH > 0.05; Figure S1). This suggests that plasticity in these traits exhibits high variation overall without consistent latitudinal patterns, likely shaped primarily by local factors such as soil moisture variation, herbivore pressure, or genotype-specific tolerance mechanisms rather than broad latitudinal patterns alone.

We recognize that latitude acts as a proxy for multiple selective pressures beyond temperature and precipitation, including photoperiod, frost frequency, and growing season length, which together shape the evolution of plasticity strategies.

### Conclusions and Implications for Conservation and Agriculture

Our investigation reveals a complex landscape of genotype-by-environment interactions in *F. vesca*. Phenotypic variation is driven primarily by the environment (29%), but a substantial portion is explained by genotype (9%) and, critically, their interaction (18%). We demonstrate that plasticity is not uniform but is itself an evolved, trait-specific characteristic.

These insights have direct applications for conservation and agricultural breeding. For assisted migration, plasticity profiles in key metabolites can guide the selection of donor populations tailored to recipient sites. Managers should screen candidate genotypes for demonstrated flexibility in stress-related metabolites rather than relying solely on geographic origin. Because metabolic adjustments generally occur rapidly but structural acclimation unfolds over longer timescales, relocation programs should include multi-year performance monitoring to capture both immediate and delayed responses before evaluating establishment success.

In crop breeding, our results caution against assuming a simple hierarchy of trait lability. Instead, breeders can leverage trait-specific plasticity markers to select alleles conferring targeted environmental responsiveness. Introgressing alleles that enhance plasticity in stress-related pathways (e.g., temperature-responsive organic acid metabolism) into cultivated strawberry may improve resilience to climate extremes without compromising developmental stability. Furthermore, understanding genotype-specific plasticity patterns can inform precision management, adjusting irrigation, shading, or photoperiod regimes to match each cultivar’s adaptive potential and thereby optimize yield and quality under variable field conditions.

## Materials and Methods

### Plant material

A total of fifteen *Fragaria vesca* genotypes were selected from the previous characterized European germplasm collection comprising approximately 200 genotypes originating from across the continent (Urrutia et al., 2023; Toivainen et al., 2024). The fifteen *F. vesca* genotypes were chosen to represent a south – north gradient in their geographic origin across Europe (Figure 1). For consistency with the broader germplasm collection, each genotype retained its original collection identifier (e.g., ES2, FIN13), though they are also referred to with the prefix ’G’ (for ’genotype’) followed by a number from 1 to 16 for this study. Genotype G08 (GER3) was excluded from this study due to insufficient replication. The numbering denoted the latitudinal origin of each genotype, increasing progressively from southern to norther locations, with detailed geographical coordinates and original collection identifiers provided in Table S1 (Figure 1, Table S1).

A total of 10 plants per genotype, considered as biological replicates, were grown in four different environments along a south–north gradient. These environments, hereafter referred to as ‘common gardenś were established in Gontrode, Belgium (50°59_′_0.581_″_N, 3°47_′_50.248_″_E); Alnarp, Sweden (55°38_′_59.99_″_N, 13°03_′_60.00_″_E); Ruissalo, Finland (60°26’0.920”N, 22°10’23.429”E); and Kevo, Finland (69°34_′_51_″_N, 026°42_′_56_″_E). Genotype G09 (LIT3) was absent from the Kevo site throughout the study due to propagation limitations. Plants were monitored over two growing seasons with contrasting weather conditions, 2021 and 2022. To account for both spatial and temporal variation, we use the term ‘GARDEN_ID’ to denote each location × year combination (e.g. Alnarp 2021 is represented as ALN_21). Due to logistical constraints, sampling was not possible in all locations across both years. Data were collected from GON_21, ALN_21 and RUI_21 in 2021, and from GON_22 and KEV_22 in 2022. In each location, plants were arranged in spatial blocks, with one plant per genotype per block, to control microenvironmental variation. All plants were grown under open-air conditions with natural temperature and precipitation. Propagation, establishment, maintenance protocols and further details on the experimental design are provided in (De-la-Cruz et al., 2025a).

## Data collection

### Plant performance-related measurements

A total of thirteen plant performance-related traits were measured, including six biomass-related traits, four reproduction-related traits, and three herbivore damage-related traits. These traits were quantified per plant per year, based on measurements taken at fixed points during the growing season: before flowering, during peak flowering and during peak fruiting. All three timepoints were present for 2021 and only the first and latter for 2022 (no peak flowering measurement).

### Biomass traits

For each plant, we measured the number of leaves, number of rosettes, and plant volume. To account for their growth form, we modelled plants as half spheroids. Accordingly, volume was calculated as one-half of the full spheroid volume, using the equation: volume = 0.5 × (4/3π × height² × width). For each of these measurements, a mean (across all timepoints) and a relative change (final minus initial value, divided by the initial value) during the season was calculated. This resulted in the following biomass and growth traits: (i) the absolute mean number of produced leaves during the whole season, (ii) the change in number of leaves, (iii) the absolute mean number of rosettes per plant, (iv) the change in number of rosettes, (v) absolute mean plant volume in cm³, (vi) the change in plant volume in cm³.

### Reproductive traits

We used measurements from both the sexual and asexual modes of reproduction: fruits and runners produced. This resulted in the following four traits: (i) the total (summed) number of ripe fruits produced by each plant throughout the season (ii) the total (summed) number of runners produced, (iii) the mean volume of produced fruits of a plant, estimated by measuring the length and width of up to 5 randomly selected fruits at each time point and considering them as spheroids, and averaging the spheroid volumes across the five fruits and the different time points and (iv) the mean proportion of fertilized achenes, calculated per plant by randomly selecting up to 5 fruits per plant, then randomly selecting 10 achenes from each of these fruits at maturity and scoring fertilization (1 = fertilized, 0 = unfertilized), then averaging across the 10 achenes per fruit and then across the selected fruits per plant.

### Herbivore damage

Herbivore damage was measured by estimating the area of the leaves consumed by herbivores on ten randomly selected leaves per plant. To estimate the overall impact of herbivores on each plant, the following three traits were calculated: (i) the mean percentage of leaf damage over the season, (ii) maximum percentage of leaf damage at any time during the season, and (iii) coefficient of temporal variation of leaf damage over the season (standard deviation divided by the mean).

### Metabolite measurements

A total of 3-5 young leaves per plant were collected and immediately frozen in liquid nitrogen to halt metabolic activity and stored at -80°C. Each plant was considered a biological replicate. Within each of the four common gardens, leaf samples were collected in July of both 2021 and 2022, except for GON_22 where sampling occurred in May to ensure a comparable developmental stage. Primary metabolite extraction and analysis by Gas Chromatography-time-of-flight-Mass Spectrometry (GC-TOF-MS) were carried out as previously reported by Vallarino et al., 2018. The resulting data, representing the abundance of 36 primary metabolites, were scaled by dry weight and relative to control samples. These control samples consisted of an arbitrary mixture of all 15 genotypes, ensuring comprehensive metabolite identification and allowing for the differentiation between the absence of a metabolite due to biological factors versus technical limitations of the GC-MS. Identified primary metabolites included thirteen amino acids, eleven organic acids, eight sugars and sugar derivatives, three polyols, and one nitrogen-related compound (urea) (Table S2).

### Data Preprocessing

Raw metabolite abundances and performance trait measurements for each individual plant were logD-transformed and then z-score scaled to ensure comparability across traits with different units and variances. For Partial Least Squares Discriminant Analysis (PLS-DA) and Permutational Multivariate Analysis of Variance (PERMANOVA), the transformed, scaled values were used directly. For hierarchical cluster analysis (HCA), Pearson correlations and heatmaps, trait means per genotype–garden–year combination were likewise logD-transformed and z-scored before analysis. This standardized workflow guarantees that all traits contribute equally to distance calculations and multivariate models.

### Data visualization and analysis

All statistical analyses were based on the full set of individual replicate values for each metabolite and trait. To explore metabolic responses to environmental variation, we performed HCA on mean-centered and scaled metabolite data alongside Pearson correlations between metabolite levels and monthly climatic variables (June–August precipitation; mean, maximum, and minimum temperature). These relationships were visualized in combined heat maps to interpret metabolic patterns relative to climate.

To assess trait variation between garden environments, genotypes, and their interaction, we applied PLS-DA and PERMANOVA, which was conducted using Euclidean distances and 9999 permutations to partition phenotypic variance among environment (GARDEN_ID), genotype, and G×E interaction.

To partition variance explained by climatic origin versus garden environment, we performed distance-based redundancy analysis (db-RDA) with sequential conditioning for four functional trait groups (metabolic, biomass, reproduction, and herbivore damage-related traits). Again, Euclidean distances and 9999 permutations were employed. Annual Mean Temperature (BIO_1) and Annual Precipitation (BIO_12) were tested as bioclimatic predictors (WorldClim v2.1; Fick and Hijmans, 2017). For each climatic variable, three models were employed using db-RDA (see Methods S1 for model details). The magnitude of phenotypic plasticity for each genotype was quantified by calculating intraclass correlation coefficients (ICCs) derived from linear mixed models. For each trait and genotype, ICCs were computed using trait values across all GARDENID environments, with ICC values ranging from 0 (highly plastic, maximum environmental sensitivity) to 1 (fully canalized, consistent across environments).

Principal Component Analysis (PCA) on ICC matrices were performed to identify general plasticity patterns among genotypes and traits. To test whether plasticity profiles were latitudinally structured, we conducted db-RDA with latitude as the explanatory variable.

Relationships between individual trait plasticity (ICC values) and latitudinal origin of each genotype were tested using Pearson correlations. P-values were adjusted for multiple testing using the Benjamini–Hochberg (BH) procedure. Significant correlations reveal which specific traits show latitude-dependent patterns of plasticity.

All analyses were conducted in R version 4.3.1. Full model specifications and detailed data structures are provided in Methods S1. Complete code and data processing pipelines are available in the public GitHub repository: https://github.com/jjoseenrique/NP_JEPM.

## Data Availability Statement

All data are available within the manuscript. The data supporting the findings are available in Figure S1; Tables S1-S4. Raw metabolomic data and performance-related traits (biomass, reproduction, and herbivore damage-related traits) will be deposited in Zenodo upon manuscript acceptance and made publicly available with a DOI.

## Acknowledgements

This work was financially supported by the European Commission (BiodivERsA project PlantCline: Adapting plant genetic diversity to climate change along a continental latitudinal gradient, project ID BiodivClim-177; PCI2020-120719-2), and the Spanish Ministry of Science and Innovation (PID2021-128527OB-I00 to SO and PID2021-123677OB-I00 to DP). The AI tools were only used for grammar corrections and minor text improvements, as well as for consistent code documentation.

## Author contributions

SO, DP and DB planned and conceptualized the research. JEP-M conducted metabolic profiling as well as all statistical analyses, data interpretation, and manuscript writing. DB and FB provided computational guidance and supervised all analytical steps, closely reviewing the methodological approach and results. MLV contributed statistical expertise and suggested analytical improvements and alternative approaches. FB, MLV, IMD-L-C, CD, and A M collected plant samples and performed performance-related traits measurements. DP and SO supervised the entire project, provided conceptual guidance throughout the study, and managed manuscript submission and revision. All authors read and approved the final manuscript.

## References

Acasuso-Rivero, C., Murren, C. J., Schlichting, C. D., & Steiner, U. K. (2019). Adaptive phenotypic plasticity for life-history and less fitness-related traits. Proceedings of the Royal Society B, 286, 0653. 10.1098/rspb.2019.0653

Ackerly, D. D. (2003). Community assembly, niche conservatism, and adaptive evolution in changing environments. International Journal of Plant Sciences, 164, S165–S184. 10.1086/368401

Agrawal, A. A. (2001). Phenotypic plasticity in the interactions and evolution of species. Science, 294(5541), 321–326. 10.1126/science.1060701

Bartels, D., & Sunkar, R. (2005). Drought and salt tolerance in plants. Critical Reviews in Plant Sciences, 24(1), 23–58. 10.1080/07352680590910410

Bongers, F. J., Olmo, M., Lopez-Iglesias, B., Anten, N. P. R., & Villar, R. (2017). Drought responses, phenotypic plasticity and survival of Mediterranean species in two different microclimatic sites. Plant Biology, 19(3), 386–395. 10.1111/plb.12544

Bradshaw, A. D. (1965). Evolutionary significance of phenotypic plasticity in plants. Advances in Genetics, 13, 115–155. 10.1016/s0065-2660(08)60048-6

Butnariu, M., & Bocso, N.-S. (2022). The biological role of primary and secondary plant metabolites. Nutrition and Food Processing, 5(4), 1–7. 10.31579/2637-8914/094

Comas, L. H., Becker, S. R., Cruz, V. M. V., Byrne, P. F., & Dierig, D. A. (2013). Root traits contributing to plant productivity under drought. Frontiers in Plant Science, 4, 442. 10.3389/fpls.2013.00442

Corpas, F. J., Gupta, D. K., & Palma, J. M. (2021). Tryptophan: A precursor of signaling molecules in higher plants. In Hormones and plant response (pp. 273–289). Springer. 10.1007/978-3-030-77477-6_11

Cranston, B. H., Monks, A., Whigham, P. A., & Dickinson, K. J. (2015). Variation and response to experimental warming in a New Zealand cushion plant species. Austral Ecology, 40(6), 642–650. 10.1111/aec.12231

De Kort, H., Panis, B., Helsen, K., Douzet, R., Janssens, S. B., & Honnay, O. (2020). Pre-adaptation to climate change through topography-driven phenotypic plastic-ity. Journal of Ecology, 108(4), 1465–1474. 10.1111/1365-2745.13365

De Lisle, S. P., & Rowe, L. (2023). Condition dependence and the paradox of missing plasticity costs. Evolution Letters, 7(1), 67–78. 10.1093/evlett/qrad009

De Villemereuil, P., Mouterde, M., Gaggiotti, O. E., & Till-Bottraud, I. (2018). Patterns of phenotypic plasticity and local adaptation in the wide elevation range of the al-pine plant Arabis alpina. Journal of Ecology, 106(5), 1952–1971. 10.1111/1365-2745.12955

De-la-Cruz, I. M., Batsleer, F., Bonte, D., Diller, C., Hytönen, T., Muola, A., et al. (2022). Evolutionary ecology of plant-arthropod interactions in light of the “omics” sciences: A broad guide. Frontiers in Plant Science, 13, 808427. 10.3389/fpls.2022.808427

De-La-Cruz, I. M., Batsleer, F., Bonte, D., Diller, C., Hytönen, T., Luis Izquierdo, J., et al. (2025a). Genotypic responses to different environments and reduced precipitation reveal signals of local adaptation and phenotypic plasticity in woodland strawberry. Annals of Botany, 136(3), 611–621. 10.1093/aob/mcaf025

De-La-Cruz, I. M., Batsleer, F., Diller, C., Izquierdo, J. L., Still, S., Osorio, S., et al. (2025b). Flowering responses of the woodland strawberry to local climate and reduced precipitation along a European latitudinal gradient. Journal of Plant Ecology, 18(5). 10.1093/jpe/rtaf105

Eckardt, N. A. (2010). Myo-Inositol biosynthesis genes in Arabidopsis: Differential patterns of gene expression and role in cell death. Plant Cell, 22(3), 537–538. 10.1105/tpc.110.220310

Edger, P. P., Poorten, T. J., VanBuren, R., Hardigan, M. A., Colle, M., McKain, M. R., et al. (2019). Origin and evolution of the octoploid strawberry genome. Nature Genetics, 51(3), 541–547. 10.1038/s41588-019-0356-4

Edger, P. P., VanBuren, R., Colle, M., Poorten, T. J., Wai, C. M., Niederhuth, C. E., et al. (2018). Single-molecule sequencing and optical mapping yields an improved ge-nome of woodland strawberry (Fragaria vesca) with chromosome-scale contiguity. GigaScience, 7(2), 1–7. 10.1093/gigascience/gix124

Fick, S. E., & Hijmans, R. J. (2017). WorldClim 2: New 1-km spatial resolution climate surfaces for global land areas. International Journal of Climatology, 37(12), 4302–4315. 10.1002/joc.5086

Flores, S., Forister, M. L., Sulbaran, H., Díaz, R., & Dyer, L. A. (2023). Extreme drought disrupts plant phenology: Insights from 35 years of cloud forest data in Venezuela. Ecology, 104(10), e4012. 10.1002/ecy.4012

Gaskell, D. E., Huber, M., O’Brien, C. L., Inglis, G. N., Acosta, R. P., Poulsen, C. J., et al. (2022). The latitudinal temperature gradient and its climate dependence as inferred from foraminiferal δ18O over the past 95 million years. Proceedings of the National Academy of Sciences of the United States of America, 119(20), e2111332119. 10.1073/pnas.2111332119

Gornish, E. S., & Tylianakis, J. M. (2013). Community shifts under climate change: mechanisms at multiple scales. American Journal of Botany, 100(7), 1422–1434. 10.3732/ajb.1300046

Granot, D., David-Schwartz, R., Kelly, G., Hellmann, H. A., Jang, J. C., Jeon, J.-S., et al. (2013). Hexose kinases and their role in sugar-sensing and plant development. Frontiers in Plant Science, 4, 44. 10.3389/fpls.2013.00044

Gratani, L. (2014). Plant phenotypic plasticity in response to environmental factors. Advances in Botany, 2014, 1–17. 10.1155/2014/208747

Gu, Z., Eils, R., & Schlesner, M. (2016). Complex heatmaps reveal patterns and correlations in multidimensional genomic data. Bioinformatics, 32(18), 2847–2849. 10.1093/bioinformatics/btw313

Heide, O. M., & Sønsteby, A. (2007). Interactions of temperature and photoperiod in the control of flowering of latitudinal and altitudinal populations of wild strawberry (Fragaria vesca). Physiologia Plantarum, 130(2), 280–289. 10.1111/j.1399-3054.2007.00906.x

Hollander, J., Snell-Rood, E., & Foster, S. (2015). New frontiers in phenotypic plasticity and evolution. Heredity, 115(4), 273–275. 10.1038/hdy.2015.64

Ibáñez, I., Gornish, E. S., Buckley, L., Debinski, D. M., Hellmann, J., Helmuth, B., et al. (2013). Moving forward in global-change ecology: capitalizing on natural variability. Ecology and Evolution, 3(1), 170–181. 10.1002/ece3.433

Jentsch, A., Kreyling, J., Boettcher-Treschkow, J., & Beierkuhnlein, C. (2009). Beyond gradual warming: extreme weather events alter flower phenology of European grassland and heath species. Global Change Biology, 15(4), 837–849. 10.1111/j.1365-2486.2008.01690.x

Joschinski, J., & Bonte, D. (2021). Diapause and bet-hedging strategies in insects: a meta-analysis of reaction norm shapes. Oikos, 130(8), 1240–1250. 10.1111/oik.08116

Joshi, S. (2023). Plant adaptations to extreme environments: Survival strategies for plants in harsh or unique habitats—A review. Advance Research in Sciences, 1(1), 1–6. 10.54026/ars.1010

Journée, M., Goudenhoofdt, E., Vannitsem, S., & Delobbe, L. (2023). Quantitative rainfall analysis of the 2021 mid-July flood event in Belgium. Hydrology and Earth System Sciences, 27(8), 3169–3189. 10.5194/hess-27-3169-2023

Karban, R., & Baldwin, I. T. (2007). Induced responses to herbivory. University of Chicago Press.

Kassambara, A., & Mundt, F. (2020). factoextra: Extract and visualize the results of multivariate data analyses. https://CRAN.R-project.org/package=factoextra

Kaur, G., & Asthir, B. (2015). Proline: A key player in plant abiotic stress tolerance. Biologia Plantarum, 59(3), 609–619. 10.1007/s10535-015-0549-3

Kawecki, T. J., & Ebert, D. (2004). Conceptual issues in local adaptation. Ecology Letters, 7(12), 1225–1241. 10.1111/j.1461-0248.2004.00684.x

Khan, N., Ali, S., Zandi, P., Mehmood, A., Ullah, S., Ikram, M., et al. (2020). Role of sugars, amino acids and organic acids in improving plant abiotic stress tolerance. Pakistan Journal of Botany, 52(2), 355–363. 10.30848/pjb2020-2(24)

Kiliç, T. (2023). Seed treatments with salicylic and succinic acid to mitigate drought stress in flowering kale cv. “Red Pigeon F1.” Scientia Horticulturae, 313, 111939. 10.1016/j.scienta.2023.111939

Körner, C., Fajardo, A., & Hiltbrunner, E. (2023). Biogeographic implications of plant stature and microclimate in cold regions. Communications Biology, 6, 1–3. 10.1038/s42003-023-05032-5

Lê, S., Josse, J., & Husson, F. (2008). FactoMineR: An R package for multivariate analysis. Journal of Statistical Software, 25(1), 1–18. 10.18637/jss.v025.i01

Li, Y., Pi, M., Gao, Q., Liu, Z., & Kang, C. (2019). Updated annotation of the wild strawberry Fragaria vesca V4 genome. Horticulture Research, 6, 15. 10.1038/s41438-019-0142-6

Liljequist, D., Elfving, B., & Roaldsen, K. S. (2019). Intraclass correlation—A discussion and demonstration of basic features. PLoS One, 14(7), e0219854. 10.1371/journal.pone.0219854

Murren, C. J., Auld, J. R., Callahan, H., Ghalambor, C. K., Handelsman, C. A., Heskel, M. A., et al. (2015). Constraints on the evolution of phenotypic plasticity: limits and costs of phenotype and plasticity. Heredity, 115(4), 293–301. 10.1038/hdy.2015.8

Neuner, G., Buchner, O., & Braun, V. (2000). Short-term changes in heat tolerance in the alpine cushion plant Silene acaulis ssp. excapa [All.] J. Braun at different alti-tudes. Plant Biology, 2(6), 677–683. 10.1055/s-2000-16635

Nievola, C. C., Carvalho, C. P., Carvalho, V., & Rodrigues, E. (2017). Rapid responses of plants to temperature changes. Temperature, 4(4), 371–405. 10.1080/23328940.2017.1377812

Oksanen, J., Simpson, G. L., Blanchet, F. G., Kindt, R., Legendre, P., Minchin, P. R., et al. (2025). Community ecology package [R package vegan version 2.6-10]. https://CRAN.R-project.org/package=vegan

Panchal, P., Miller, A. J., & Giri, J. (2021). Organic acids: Versatile stress-response roles in plants. Journal of Experimental Botany, 72(14), 4038–4052. 10.1093/jxb/erab019

Planchet, E., & Limami, A. M. (2015). Amino acid synthesis under abiotic stress. In Amino acids in higher plants (pp. 262–276). 10.1079/9781780642635.0262

Plaxton, W. C., & Tran, H. T. (2011). Metabolic adaptations of phosphate-starved plants. Plant Physiology, 156(3), 1006–1015. 10.1104/pp.111.175281

Reger, J., Lind, M. I., Robinson, M. R., & Beckerman, A. P. (2017). Predation drives local adaptation of phenotypic plasticity. Nature Ecology & Evolution, 2, 100–107. 10.1038/s41559-017-0373-6

Rohart, F., Gautier, B., Singh, A., & Lê Cao, K. A. (2017). mixOmics: An R package for ’omics feature selection and multiple data integration. PLoS Computational Bi-ology, 13(11), e1005752. 10.1371/journal.pcbi.1005752

Ruan, Y. L., Patrick, J. W., Shabala, S., & Slewinski, T. L. (2013). Uptake and regulation of resource allocation for optimal plant performance and adaptation to stress. Frontiers in Plant Science, 4, 71675. 10.3389/fpls.2013.00455

Sakamoto, A., & Murata, N. (2002). The role of glycine betaine in the protection of plants from stress: clues from transgenic plants. Plant, Cell & Environment, 25(2), 163–171. 10.1046/j.0016-8025.2001.00790.x

Schlichting, C. D. (1986). The evolution of phenotypic plasticity in plants. Annual Review of Ecology and Systematics, 17, 667–693. 10.1146/annurev.es.17.110186.003315

Schneider, H. M. (2022). Characterization, costs, cues and future perspectives of phenotypic plasticity. Annals of Botany, 130(2), 131–148. 10.1093/aob/mcac087

Shulaev, V., Sargent, D. J., Crowhurst, R. N., Mockler, T. C., Folkerts, O., Delcher, A. L., et al. (2010). The genome of woodland strawberry (Fragaria vesca). Nature Genetics, 43(2), 109–116. 10.1038/ng.740

Snell-Rood, E. C., & Ehlman, S. M. (2021). Ecology and evolution of plasticity. In Phenotypic plasticity & evolution (pp. 139–160). 10.1201/9780429343001-8

Sparks, A. H. (2018). nasapower: A NASA POWER global meteorology, surface solar energy and climatology data client for R. Journal of Open Source Software, 3(23), 1035. 10.21105/joss.01035

Tahjib-Ul-arif, M., Zahan, M. I., Karim, M. M., Imran, S., Hunter, C. T., Islam, M. S., et al. (2021). Citric acid-mediated abiotic stress tolerance in plants. International Journal of Molecular Sciences, 22, 37235. 10.3390/ijms22137235

Takahashi, K. H. (2019). Multiple modes of canalization: Links between genetic, environmental canalizations and developmental stability, and their trait-specificity. Seminars in Cell & Developmental Biology, 88, 14–20. 10.1016/j.semcdb.2018.05.018

Toivainen, T., Salonen, J. S., Kirshner, J., Lembinen, S., Kort, H. De, Lyyski, A., et al. (2024). Late Quaternary climatic impact on the woodland strawberry genome: a perennial herb’s tale. bioRxiv, 2024.10.09.617376. 10.1101/2024.10.09.617376

Traill, L. W., Lim, M. L. M., Sodhi, N. S., & Bradshaw, C. J. A. (2010). Mechanisms driving change: altered species interactions and ecosystem function through glob-al warming. Journal of Animal Ecology, 79(4), 937–947. 10.1111/j.1365-2656.2010.01695.x

Urrutia, M., Meco, V., Rambla, J. L., Martín-Pizarro, C., Pillet, J., Andrés, J., et al. (2023). Diversity of the volatilome and the fruit size and shape in European woodland strawberry (Fragaria vesca). The Plant Journal, 116(5), 1201–1217. 10.1111/tpj.16404

Valladares, F., Gianoli, E., & Gómez, J. M. (2007). Ecological limits to plant phenotypic plasticity. New Phytologist, 176(4), 749–763. 10.1111/j.1469-8137.2007.02275.x

Vallarino, J. G., de Abreu e Lima, F., Soria, C., Tong, H., Pott, D. M., Willmitzer, L., et al. (2018). Genetic diversity of strawberry germplasm using metabolomic biomarkers. Scientific Reports, 8(1), 1–13. 10.1038/s41598-018-32212-9

Vergeer, P., & Kunin, W. E. (2013). Adaptation at range margins: common garden trials and the performance of Arabidopsis lyrata across its northwestern European range. New Phytologist, 197(4), 989–1001. 10.1111/nph.12060

Via, S., & Lande, R. (1985). Genotype-environment interaction and the evolution of phenotypic plasticity. Evolution, 39(3), 505–522. 10.1111/j.1558-5646.1985.tb00391.x

Wang, L., Cui, D., Zhao, X., & He, M. (2017). The important role of the citric acid cycle in plants. Genomics & Applied Biology, 8(4), 1–5. 10.5376/gab.2017.08.0004

Weiskopf, S. R., Rubenstein, M. A., Crozier, L. G., Gaichas, S., Griffis, R., Halofsky, J. E., et al. (2020). Climate change effects on biodiversity, ecosystems, eco-system services, and natural resource management in the United States. Science of the Total Environment, 733, 137782. 10.1016/j.scitotenv.2020.137782

Wenzel, S., Cox, P. M., Eyring, V., & Friedlingstein, P. (2016). Projected land photosynthesis constrained by changes in the seasonal cycle of atmospheric CO2. Nature, 538(7625), 499–501. 10.1038/nature19772

Zhou, Y., Xiong, J., Shu, Z., Dong, C., Gu, T., Sun, P., et al. (2023). The telomere-to-telomere genome of Fragaria vesca reveals the genomic evolution of Fragaria and the origin of cultivated octoploid strawberry. Horticulture Research, 10, uhado27. 10.1093/hr/uhad027

