## Supplementary Materials for "Variation in phenotypic plasticity of metabolic and performance traits along a latitudinal gradient in Woodland Strawberry"

The following Supplementary Material is available for this article:

**Figure S1:** Correlation scatterplots between latitudinal origin and Intraclass Correlation Coefficients (ICCs) for all 49 traits measured in *Fragaria vesca* genotypes. Each point represents a genotype (n = 15), with latitude (°N) on the x-axis and trait-specific plasticity (ICC value) on the y-axis. Linear regression lines (black dashed) and 95% confidence intervals (shaded gray) are shown. Benjamini-Hochberg adjusted p-values (p-valueBH) and Pearson correlation coefficients (r) are displayed for each trait. Traits are organized by functional category: metabolic compounds (metabolites 1–36) and performance-related traits (phenotypes 1–13). Significant associations (p-valueBH < 0.05) are marked with asterisks and highlighted in color. This figure complements Figure 6 by providing individual trait-level detail of latitudinal structuring in plasticity profiles.

*Figure S1 is attached as .pdf file due to its size*

**Table S1:** Geographic origins of the fifteen Fragaria vesca genotypes, showing latitude and longitude for each collection site.

| **Genotype** | **Identifier** | **Latitude** | **Longitude** |
| --- | --- | --- | --- |
| G01 | ES2 | 37.7796° N | 3.7849° O |
| G02 | ES20 | 40.2938° N | 5.0091° O |
| G03 | ES13 | 43.1344° N | 4.8880° O |
| G04 | IT3 | 45.9400° N | 10.8100° E |
| G05 | GER100 | 47.9667° N | 7.8333° E |
| G06 | FR2 | 48.8014° N | 2.1301° E |
| G07 | FR3 | 50.0157° N | 2.6974° E |
| G09 | LIT3 | 54.5729° N | 24.6722° E |
| G10 | DK1 | 55.5703° N | 9.7466° E |
| G11 | SE6 | 60.1018° N | 18.3433° E |
| G12 | FIN13 | 60.2292° N | 25.0233° E |
| G13 | FIN39 | 60.4051° N | 25.1562° E |
| G14 | NOR21 | 70.1671° N | 24.7561° E |
| G15 | NOR19 | 70.0301° N | 22.0653° E |
| G16 | NOR8 | 70.0324° N | 23.4012° E |

**Table S2:** Raw metabolomic data for all 36 primary metabolites detected in *Fragaria vesca* leaf tissue across 581 samples grown in five common garden environments. Raw data deposited in Zenodo (DOI: xxxxx).

*Table S2 is attached as .xlsx file due to its size*

**Table S3:** Raw phenotypic data for all 13 plant performance-related traits measured in *Fragaria vesca* across 581 samples grown in five common garden environments. Traits include morphological (leaf number, rosette number, plant volume), reproductive (ripe fruits, runners, fruit volume, fertilized seeds), and herbivory-related measurements (leaf damage %). Raw data deposited in Zenodo (DOI: xxxxx).

*Table S3 is attached as .xlsx file due to its size*

**Table S4:** Mapping of trait numbers used in PCA biplots (Figure 5) to full trait names. Each trait measured in the study is assigned a unique number for visual clarity in the PCA plots, with this table providing the correspondence between trait numbers and their complete descriptive names.

| **Number ID** | **Trait Name** | **Number ID** | **Trait Name** |
| --- | --- | --- | --- |
| **1** | Pyruvic acid | **26** | Trehalose |
| **2** | Valine | **27** | Maltitol |
| **3** | Isoleucine | **28** | Galactinol |
| **4** | Glycine | **29** | Quinic acid |
| **5** | Phosphoric acid | **30** | Fructose |
| **6** | Proline | **31** | Glucose |
| **7** | Urea | **32** | Citric acid |
| **8** | Glyceric acid | **33** | Methyl α-D-glucopyranoside |
| **9** | Alanine | **34** | Dehydroascorbic acid |
| **10** | Serine | **35** | myo-Inositol |
| **11** | Succinic acid | **36** | Sucrose |
| **12** | Threonine | **37** | Mean leaf number |
| **13** | Fumaric acid | **38** | Leaf number change |
| **14** | Nicotinic acid | **39** | Mean rosette number |
| **15** | Erythritol | **40** | Rosette number change |
| **16** | Malic acid | **41** | Mean plant volume |
| **17** | GABA | **42** | Plant volume change |
| **18** | Aspartic acid | **43** | Total ripe fruits |
| **19** | Threonic acid | **44** | Total runners |
| **20** | Xylose | **45** | Mean fruit volume |
| **21** | Xylitol | **46** | Fertilized achenes (%) |
| **22** | Pyroglutamic acid | **47** | Mean leaf damage (%) |
| **23** | Glutamic acid | **48** | Leaf damage (CV) |
| **24** | Arginine | **49** | Max leaf damage (%) |
| **25** | Tryptophan |  |  |

**Methods S1:** Comprehensive description of statistical methods including correlation analyses, multivariate approaches (PLS-DA, PERMANOVA, db-RDA), quantification of phenotypic plasticity via linear mixed models and Intraclass Correlation Coefficients (ICCs), and variance partitioning by climatic origin. All analyses performed in R v4.4.0 using vegan, lme4, mixOmics, and FactoMineR packages.

Hierarchical Clustering and Pearson Correlation

To link trait expression with local climate, genotype-specific trait means were calculated for each garden–year combination (e.g., mean value of genotype G01 in GON_21). Monthly climate data (precipitation, mean/max/min temperature) for each GARDEN_ID were retrieved via the get_power function in the nasapower R package (Sparks, 2018). For metabolic traits, means were correlated with climate in the month of sampling; for performance traits, means were correlated with annual climate averages. Although Pearson correlations assume independent observations, we present them here solely as descriptive measures of trait–environment associations (N = 75 genotype–garden points). Correlation coefficients were visualized in a heatmap using the ComplexHeatmap package (Gu et al., 2016).

Analysis of Variance and Clustering (PLS-DA):

Individual plant measurements (log₂-transformed and z-scored) for all metabolites and performance traits were used. Samples were labeled by garden–year (e.g., “GON_21,” “KEV_22”), mean-centered, and unit‐variance scaled. PLS-DA was run with the plsda function from mixOmics (Rohart et al., 2017) retaining three components. Sample scores on LV1 vs. LV2 and LV1 vs. LV3 were plotted with ggplot2, adding 95% normal‐based ellipses per garden–year. Biplots were combined side by side with patchwork, consolidating legends at the bottom.

Permutational Multivariate Analysis of Variance (PERMANOVA):

PERMANOVA was performed to assess if traits are significantly different across GARDEN_IDs and/or genotypes. This analysis was conducted using the adonis2 function from vegan R package (Oksanen et al., 2025). Euclidean distances were calculated on log₂-transformed, z-scored values. Models tested GARDEN_ID, genotype, and their interaction (Type I SS), with 9999 permutations. To verify robustness, an alternative model with reversed term order (genotype first, then GARDEN_ID) was also conducted. Reported R² and p-values reflect variance explained by each term. This analysis partitions phenotypic variance among experimental factors but does not distinguish between immediate environment effects and heritable adaptation signals.

Partitioning Variance by Climatic Origin (db-RDA with Sequential Conditioning):

Distance-based redundancy analysis (db-RDA) with sequential conditioning was used to quantify variance explained by climatic origin versus garden environment for four functional trait groups (metabolic, biomass, reproduction, and herbivore damage-related traits). Euclidean distances were calculated on log₂-transformed, z-scored trait values and used as response variables in constrained ordination models using the capscale () function from the vegan R package (Oksanen et al., 2025) with 9999 permutations.

Two climatic predictors were tested separately: Annual Mean Temperature (BIO_1) and Annual Precipitation (BIO_12), obtained from WorldClim v2.1 at 2.5 arc-minute resolution (Fick and Hijmans, 2017). For each trait group and bioclimatic variable, three models were constructed:

1. GARDEN_ID model: capscale (traits ~ GARDEN_ID, distance = "euclidean") to quantify variance explained by immediate growing environment.
2. Conditional bioclimatic model: capscale (traits ~ BIO + Condition (GARDEN_ID), distance = "euclidean") to isolate variance explained by the bioclimatic variable after removing GARDEN_ID effects.
3. Full model: capscale (traits ~ GARDEN_ID + BIO, distance = "euclidean") to verify total variance explained.

Adjusted R² values were extracted using RsquareAdj() to account for the number of predictors. Statistical significance was assessed via permutation tests (9999 permutations), with p-values < 0.0001 considered highly significant.

To confirm the robustness of detected effects, marginal PERMANOVA tests were also performed using adonis2 () with by = "margin", which evaluates each predictor's independent effect while accounting for other terms in the model.

Quantification of Plasticity (Mixed Models and Intraclass Correlation Coefficients - ICC):

To quantify the plastic response of each genotype individually across the different GARDEN_ID environments, we calculated Intraclass Correlation Coefficients (ICCs). To achieve this, we performed a separate analysis for every genotype. Within each genotype's dataset, we fitted a linear mixed model for each trait using the lmer function from the lme4 package.

The model was structured with the trait value as the response variable and GARDEN_ID as a random effect (Trait ~ 1 + (1|GARDEN_ID)). The ICC was then calculated as the proportion of the total variance in the model that was attributable to the random effect (i.e., the variance among GARDEN_IDs). This procedure resulted in a matrix of ICC values, where each value represents the specific plasticity of a given genotype for a given trait. This approach allowed us to determine the extent to which variation in each trait was due to environmental factors for each genetic line separately, in line with established methods for quantifying plasticity (Liljequist et al., 2019).

Dimensionality Reduction and Plasticity Patterns (PCA):

To identify patterns in plasticity within and among GARDEN_IDs and genotypes, Principal Component Analysis (PCA) on the ICC results was performed using PCA function from the FactoMineR package (Lê et al., 2008). fviz_pca_biplot function from FactoExtra (Kassambara and Mundt, 2020) was employed for plotting, which allowed us to visualize the variance explained by each principal component and simplify the interpretation of the underlying structure of plasticity data.

Testing for a Latitudinal Gradient in Plasticity Profiles:

To formally test whether the overall plasticity profile of a genotype was structured by its geographic origin, we performed a distance-based Redundancy Analysis (db-RDA) on the full matrix of Intraclass Correlation Coefficients (ICCs) using the capscale() function from the vegan R package (Oksanen et al., 2025). ICCs were calculated for each trait within each genotype using linear mixed models with GARDEN_ID as a random effect, quantifying the consistency of trait expression across growing environments for each genotype. Euclidean distance matrices were calculated from the ICC values using vegdist() and used as the response variable in the ordination model. Latitude of origin was the sole continuous explanatory variable. The significance of the db-RDA model was assessed using an ANOVA-like permutation test (anova.cca() function) with 9,999 permutations. The proportion of variance explained in plasticity profiles by latitude was quantified using adjusted R-squared values.

Relationship Between Plasticity and Latitude (Linear Regressions):

To assess how geographic variation influences trait expression, linear regressions between the ICC values of each trait and the original latitude of the genotypes were performed using the lm function from the stats package. Model assumptions (linear relationship, homoscedasticity, normality of residuals, and independence) were evaluated via diagnostic plots. The significance and strength of these relationships were analyzed to identify the traits most sensitive to latitudinal gradients. Benjamini–Hochberg correction was applied across the list of tests (one regression per trait) to control the false discovery rate when interpreting adjusted p-values.
